# Evolutionary Influences on Local Patterns of Genetic Relatedness

**DOI:** 10.1101/2025.05.02.651970

**Authors:** T. Quinn Smith, Amatur Rahman, Stephen W. Schaeffer, Zachary A. Szpiech

**Affiliations:** Department of Biology, The Pennsylvania State University, University Park, PA 16802

## Abstract

Dimensionality reduction methods, such as Principal Components Analysis (PCA) or Multi-dimensional Scaling (MDS), when applied to genomic data, help to visualize the relatedness of individuals in lower dimensional space and are ubiquitous in population-genetic studies. These analyses use genome-wide patterns of variation to provide an “average” picture of genetic structure and relatedness. However, evolutionary processes result in different patterns of relationship among samples in local genomic regions as compared to the genome-wide aggregate. Recently, these local patterns of relatedness have been used to identify regions under selection and inverted segments. Here, we propose a unifying method to dissect these local deviations in genetic relatedness. Our method, Local Decomposition and Similarity to All Regions (LODESTAR), uses Procrustes Analysis to assess the similarity between local MDS results computed using pairwise allele sharing distances or a set of user-defined points. Given two sets of points, Procrustes Analysis computes the optimal rotation and scaling that fits one set of points onto the other, while maintaining the relative relationship between points within both sets. We use the Procrustes statistic to measure the similarity between the two sets of points. We show how this method can be used to explore local relatedness patterns that mirror sampling geography or population stratification by performing Procrustes analysis between local relatedness plots and coordinates representing sampling geography or between local relatedness plots and the genome-wide relatedness plot, respectively. In addition, we show how the variance of samples in lower-dimensional space can capture regions lacking population structure and inverted segments.

## 1 Introduction

Population structure arises from the synergistic effects of geographic, environmental, and behavioral forces subdividing a natural population into groups that evolve separately [1]. The study of how these forces create population structure is of fundamental importance to evolutionary biology; it informs our understanding of genetic relatedness and provides valuable insights into the evolutionary history of populations. A high-resolution view of relatedness will contribute to the conversation on how to best manage funds and resources to combat the endangerment of species by providing information to conservationists and evolutionary biologists about the migratory histories, gene flow, and adaptation of species.

The classical measure of population structure is Wright’s *F*_*ST*_, which measures the degree of genetic separation between two populations [2]. *F*_*ST*_ is sufficient for populations with a simplistic demographic history but fails to properly describe structure for populations with complex demographic histories [3]. Furthermore, *F*_*ST*_ requires the accurate assignment of samples to their population of origin and cannot be used to compare structure between multiple populations [4]. The limitations of *F*_*ST*_ motivated the development of more robust tools to investigate population structure.

Dimensionality reduction techniques, such as principal component analysis (PCA) and multidimensional scaling (MDS) mitigate the shortcomings of measuring population structure with *F*_*ST*_ by providing continuous axes of genetic variation [5]. Lower dimensional representations of genetic relatedness offer insight into the genealogical history between samples and provide a high-resolution view of population stratification [6, 7]. In addition, dimensionality reduction techniques are known to recapitulate sampling geography and reflect variation in cultural factors, such as linguistics [8, 9, 10, 11].

Analyses that use dimensionality reduction techniques to investigate population structure consider genome-wide markers in aggregate to summarize the relatedness between samples. However, evolutionary processes can create local genomic patterns of relatedness among samples that substantially differ from the genome-wide aggregate and would be missed in a typical genome-wide PCA or MDS analysis [12]. Several existing methods successfully utilize dimensionality reduction techniques within local genomic regions to identify targets of positive selection [13, 14]. More recently, flexible methods have been developed to visualize variation of relatedness patterns within reduced dimensions along genomic segments [12, 15, 16]. Such methods show distinct patterns of variation associated with inverted regions [15] and regions under positive selection [12].

Currently, methods calculate relatedness between samples in local genomic regions, and then, visualize the relatedness between windows in a lower-dimensional space [12]. The nested use of dimensionality reduction and aggregate view of relatedness mutes potentially informative signals in individual windows. Furthermore, geographic distributions are believed to provide insight into these evolutionary forces, which ultimately drive adaptation and genetic diversity [17]. Our goal is to explore the local genomic regions associated with such evolutionary forces. We provide a unified framework to measure the correlation between local patterns of genetic relatedness and another set of points. We introduce LOcal DEcomposition and Similarity to All Regions (LODESTAR). LODESTAR realizes local patterns of relatedness and the genome-wide pattern of relatedness by computing MDS plots using markers in local genomic regions and genome-wide markers, respectively. The goal of identifying genomic regions that are structurally similar to another plot of relatedness can be re-casted as performing Procrustes analysis between each set of points. Procrustes analysis computes the optimal rotation and scaling that fits one set of points onto the other, while maintaining the relative relationship between points in both sets [18]. The Procrustes statistic is used to measure the correlation, or equivalently similarity, between the two sets of points by summarizing the distance between corresponding points in each set.

We intend for LODESTAR to be used as an exploratory tool to inspect genomic regions with potential evolutionary significance. First, we show LODESTAR’s ability to provide candidate genomic regions that mirror the geographic relatedness of samples in the 1000 Genomes Project [19] and the 1001 Genomes Project [20]. Then, we use LODESTAR to highlight genomic regions where population stratification is preserved in both datasets. Next, we show how LODESTAR can be used to signal regions devoid of population structure in 1000 Genomes Project populations. Finally, we show how inverted segments show distinct patterns of relatedness, using *Drosophila pseudoobscura* as an example [21]. We conclude with a discussion of LODESTAR’s limitations and the broader interpretation of LODESTAR’s results.

## 2 Materials and Methods

LODESTAR’s first phase consists of calculating the relatedness of samples in a lower-dimensional space. The genome is divided into, possibly overlapping, windows. The genetic distance between the samples is calculated within each window, and cMDS is used to represent the proximity of the samples in lower-dimensional space. Simultaneously, the genome-wide relatedness plot is calculated using all haplotypes. The second phase consists of performing Procrustes analysis between each of the local plots of relatedness and the genome-wide aggregate plot of relatedness, or a user-defined set of coordinates.

### 2.1 Relatedness along a sliding window

Consider *N* diploid samples, each with phased genotypes on an *L* locus chromosome. We define a sliding window along our chromosome using parameters *H, W*, and *S*, where *H* is the size of a haplotype in number of loci, *W* is the window size in number of haplotypes, and *S* ≤ *W* is the step size in number of haplotypes, respectively. The genome is partitioned into 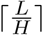 non-overlapping haplotypes. We use the partitioned genome to create a sequence of 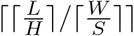, possibly overlapping, windows, such that the *k*^*th*^ window, *w*^*k*^, contains the interval of haplotypes from the *S*(*k* − 1) + 1 haplotype to the *S*(*k* − 1) + *W* haplotype, inclusively. Here, ⌈*x*⌉ denotes the ceiling of a real number *x*.

We denote the number of identical-by-state (IBS) haplotypes between samples *i* and *j* at the *h*^*th*^ haplotype as 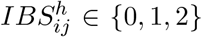. IBS is used as a measure of genetic similarity to capture the population stratification between samples [22, 23]. The genetic similarity between samples *i* and *j* in the *k*^*th*^ window is calculated as

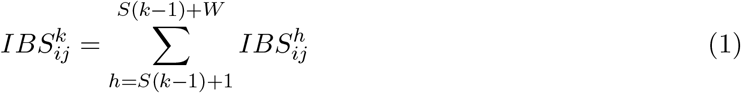

From Equation 1, we define the allele-sharing distance between samples *i* and *j* in the *k*^*th*^ window as

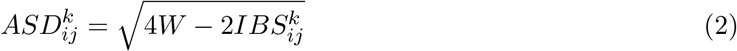

Computing Equation 2 for all pairs of samples, we form an *N* × *N* distance matrix, *ASD*^*k*^. The relatedness between samples is visualized by reducing the dimension of *ASD*^*k*^ using cMDS. The Equation 2 is a nonlinear transformation of Equation 1 such that Equation 2 is Euclidean, which allows for the use of cMDS [24]. Given a dimension *M < N*, cMDS projects the *N* samples into *M*-dimensional space while maintaining the pairwise Euclidean distance between samples. We call the lower-dimensional representation of the samples within the *k*^*th*^ window *X*_*k*_. Since *X*_*k*_ is mean-centered, 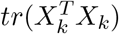 provides a measure of variance across the *M* features. The standard deviation of *X*_*k*_ is more numerically manageable, and we refer to it as 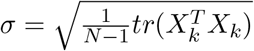.

### 2.2 Procrustes Analysis

Procrustes analysis is used to compare the overall structure between two sets of points by computing the optimal rotation, scaling, and translation of the first set of points such that the squared residuals between the corresponding points in both sets is minimized [24]. Consider two *N* × *M* matrices, *X* and *Y*. Let *x*_0_ and *y*_0_ be the centroids of *X* and *Y*, and let *x*_*r*_ and *y*_*r*_ refer to the *r*^*th*^ row of *X* and *Y*, and let *X*_*c*_ and *Y*_*c*_ be the mean-centered and normalized matrices of *X* and *Y*, respectively. Procrustes analysis creates the function *f* (*x*_*r*_) = *ρA*^*T*^ *x*_*r*_ + *b* such that

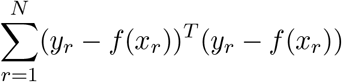

is minimized [18]. Here, *A* represents the optimal rotation, *ρ* represents the optimal scaling, and *b* represents the optimal translation. The three values, *A, ρ*, and *b*, can be found from the following singular value decomposition of 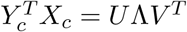.

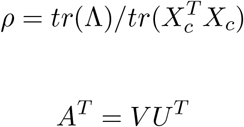

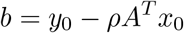

The sum of squared residuals, *ss*, is calculated by

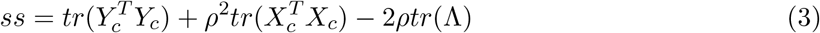

From Equation 3, we form the Procrustes statistic, 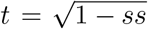, which is equivalent to the Procrustes correlation in [25]. The Procrustes statistic summarizes the structural similarity between *X* and *Y*. There is a high similarity between *X* and *Y* when *t* is close to 1, and conversely, high dissimilarity when *t* is close to 0.

### 2.3 Implementation

The user is free to define the haplotype size *H*, the window size *W*, and the step size *S*, along with the reduced dimension *M < N* given a set of *N* samples. For each window, there are *W* haplotypes and 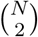 pairs of samples, resulting in *O*(*WN* ^2^) IBS calculations. All monomorphic sites are excluded from the distance calculation since they do not contribute to genetic differences between samples. The lower dimensional plot of relatedness is computed in *O*(*MN* ^2^) for small *M*, as opposed to cMDS’s traditional runtime of *O*(*N* ^3^), due to LAPACK’s *dsyevr* routine, making the complexity of each window *O*(*WN* ^2^) [26]. For haplotypes with missing data, pairwise comparisons for the haplotype are omitted from the distance calculation. For windows with much missing data or few loci, the *ASD* matrix may be of low rank, causing *dsyevr* to not converge. In such as case, the window is dropped from further analysis. We favor the combined use of ASD and cMDS as opposed to performing PCA on the genotype matrix for several reasons. The first is the felxibility to accommodate a wide variety of genotype data, and the second is to accommodate missing genotypes. In either situation, PCA on the genotype matrix is restricted to biallelic markers that must be omitted if a missing allele is present or imputed, which is error-prone, especially in the presence of low-coverage data or in the absence of a high-quality reference genome [27]. Local relatedness plots reported by LODESTAR are the Procrustes transformed coordinates. LODESTAR has the option to parallelize the distance calculations and cMDS computations across multiple threads, making the analysis feasible for thousands of samples and windows. By default, Procrustes analysis is performed between each window’s relatedness plot and the genome-wide relatedness plot. The user can supply a custom plot of points to perform Procrustes analysis against, such as a map of sampling locations. A schematic of the LODESTAR analysis is shown in Figure 1.

**Figure 1:**
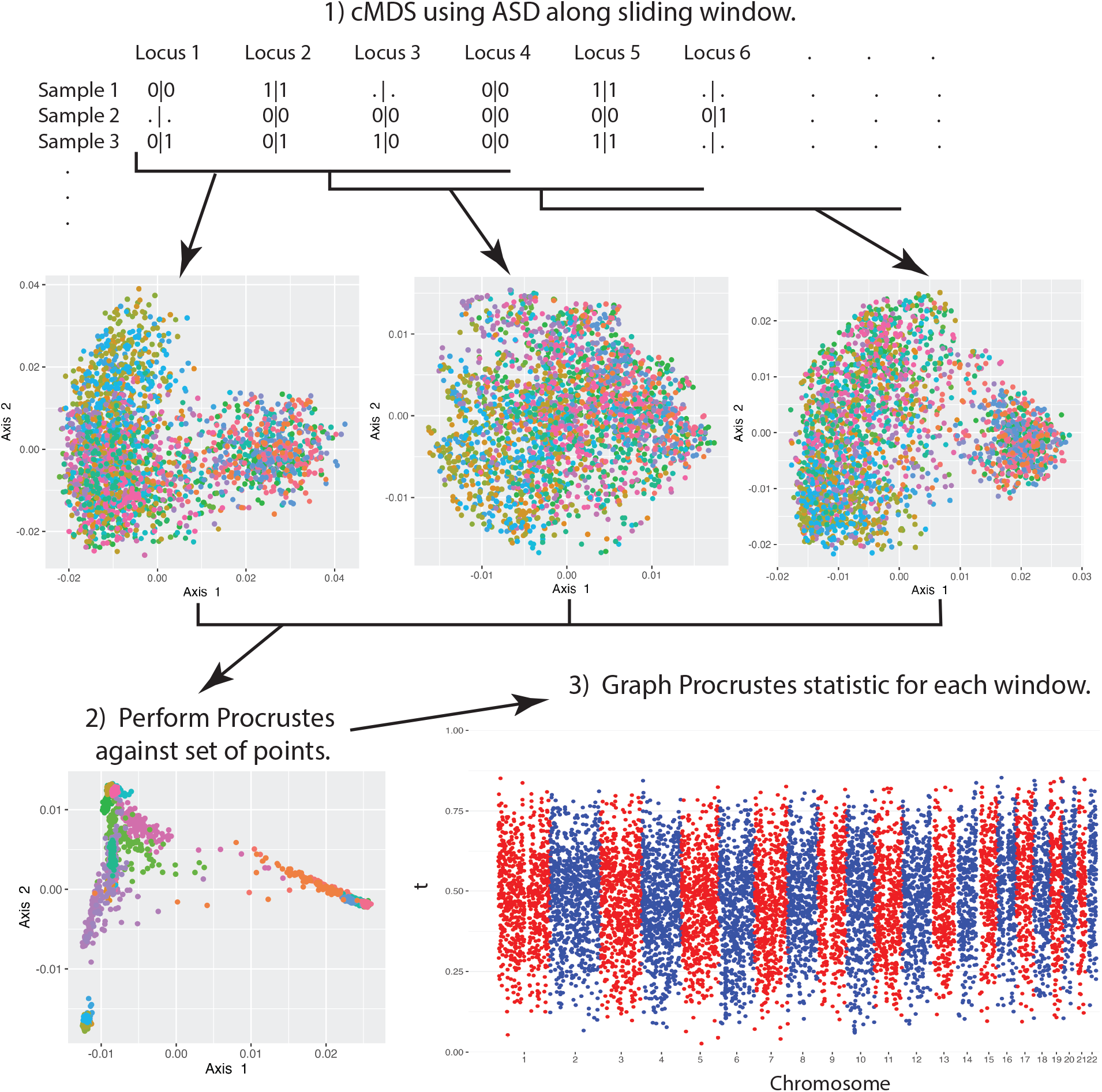
LODESTAR analysis. A schematic of the LODESTAR method. cMDS, classical multidimensional scaling. ASD, allele-sharing distance.

### 2.4 Data Sets and Preprocessing

#### Human

We defined a group of European individuals from the 1000 Genomes Project [19] composed of 107 Iberian individuals from Spain (IBS), 91 British individuals from Great Britain and Scotland (GBR), 99 Finish individuals from Finland (FIN), and 107 Tuscan individuals from Italy (TSI). We omitted the Northern Europeans from Utah (CEU) population, since their country of origin is ambiguous. We acquired the geographic centroids for the four countries. We transformed the centroids, given the latitude and longitude, (*ϕ, λ*), to the Cartesian plane using the transformation (*Rcos*(*ϕ*)*cos*(*λ*), *Rcos*(*ϕ*)*sin*(*λ*)) where *R* = 6371, the radius of the Earth in kilometers. All genomic coordinates are according to the GRCh38 reference genome. All multialleic sites were combined into one VCF record.

##### A. thaliana

We considered 919 strains of *A. thaliana* from the 1001 Genomes Project [20]. The 919 strains were taken from 14 populations (Table S2). From the 1001 Genomes accession information, we obtained the sampling location of each strain and transformed the geographic location to the Cartesian plane. Given the latitude and longitude, (*ϕ, λ*), of the sampling location in radians, we formed the Cartesian coordinates with (*Rcos*(*ϕ*)*cos*(*λ*), *Rcos*(*ϕ*)*sin*(*λ*)) where *R* = 6371, the radius of the Earth in kilometers. For each population, we formed the population’s Cartesian centroid by averaging the Cartesian sampling locations of all samples in the population. All genomic coordinates are according to the TAIR10 reference [28].

##### D. pseudoobscura

Dobzhansky and Sturtevant discovered that the third chromosome in *D. pseudoobscura* populations in western North America were segregating for over 30 different gene arrangements that were generated through overlapping paracentric inversion mutations [29]. This polymorphism was discovered using the staining of polytene salivary chromosomes developed by Painter [30]. One gene arrangement was chosen as the arbitrary Standard arrangement. New gene arrangements differing from the Standard chromosome were named for the locations where they were first collected. We considered the 8 strains carrying the Standard (ST) arrangement and the 14 strains carrying the Arrowhead (AR) arrangement. The ST and AR arrangement differs by one inverted segment. Fuller et al. also mapped the breakpoints for the sequenced gene arrangements and found that proximal and distal regions were less differentiated than central inverted regions of the chromosomes providing an ideal dataset as a test case for LODESTAR [21]. All genomic coordinates are according to the *D. pseudoobscura* reference strain FLYBASE version 3.02, which contains the AR rearrangement [31].

#### Preprocessing

LODESTAR has several preprocessing options. Sites with more than 64 alternative alleles are dropped. Unless otherwise specified, biallelic records with a minor allele frequency below 5% are excluded, record with more than 10% missing data are excluded, and windows that span more than 1 MB are dropped from further analysis. By default, cMDS is performed in *M* = 2.

### 2.5 Data Availability

LODESTAR is implemented in the C programming language. All source code and precompiled binaries are available at https://github.com/TQ-Smith/LODESTAR, along with the R script used to create graphs from LODESTAR’s output. The user manual with examples can be found at https://github.com/TQ-Smith/LODESTAR/wiki.

The data sets used can be accessed accordingly: 1000 Genomes Project Phase 3 http://ftp.1000genomes.ebi.ac.uk/vol1/ftp/release/20130502/ [19], *Drosophilia pseudoobscura* https://scholarsphere.psu.edu/resources/f488eb1f-bdf2-4497-87c0-95974fa575f2 [21], and 1001 Genomes v3 https://1001genomes.org/data/GMI-MPI/releases/v3.1/ [32]. Geographic information of the centroids of countries from the 1000 Genomes Project was accessed from https://github.com/gavinr/world-countries-centroids [33]. Geographic and population information for the 1001 Genomes was taken from https://1001genomes.org/accessions.html.

## 3 Results

### 3.1 Genes that Mirror Geography

Several studies have shown that the PCA plots of individuals using genome-wide biallelic markers reflects the geographic relatedness between the samples’ country of origin [8, 34]. Our goal was to propose candidate genomic regions such that the genetic relatedness between samples within the proposed regions reflects the geographic relatedness of the samples’ country of origin. The POPRES dataset used by Novembre et al. is too sparse to locate distinct regions [35]. We focused on a set of European individuals from the 1000 Genomes Project [19]. For our LODESTAR analysis, we compared the local plot of relatedness to the centroid of the samples’ country of origin in the Cartesian plane. The centered and scaled relatedness plot used for comparison in Procrustes analysis (Figure 2c). Local plots of relatedness were calculated in non-overlapping 1000 loci windows (Figure 2a).

**Figure 2:**
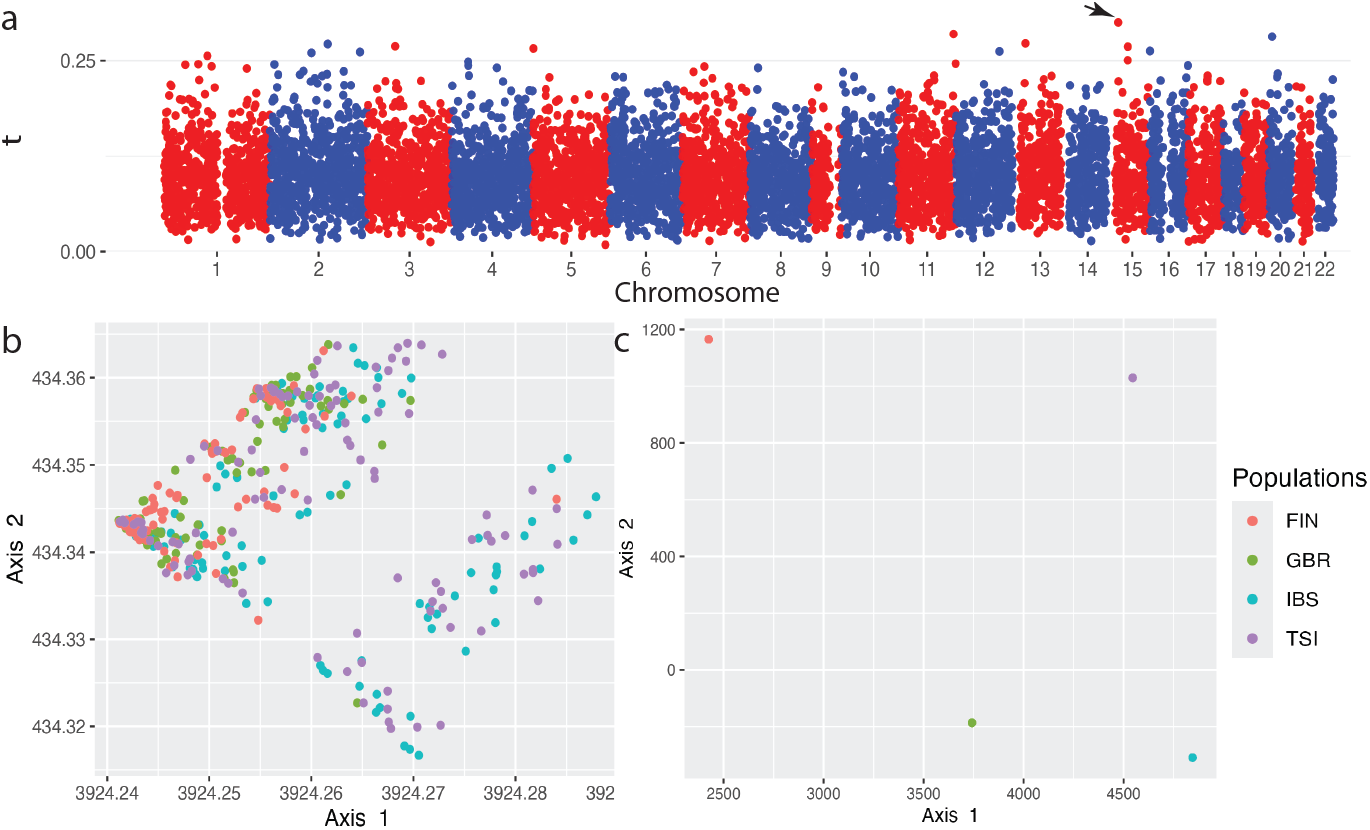
Europe Geographic Similarity. **Panel a** The Procrustes statistic plotted along the European genomes. Procrustes analysis was performed using non-overlapping, 1000 loci windows. The window indicated by the arrow is shown in Panel b. **Panel b** The relatedness within the window spanning *chr*15 : 27915418 − 28439434 is the window with the highest Procrustes statistic in the genome of *t* = 0.30. **Panel c** The centered and scaled plot of the centroids of the four countries in the Cartesian plane.

The window (Figure 2b) covering the region *chr*15 : 27915418 − 28439434 has the highest *t* = 0.30. We repeated the analysis using non-overlapping 100 ten loci haplotype windows. We found that the same window defined by *chr*15 : 27915418 − 28439434 has the highest *t*-statistic. The use of haplotypes increased the *t*-statistic to *t* = 0.354. The window defined by the region (Figure 2b) overlaps the *OCA*2 gene and completely contains the *HERC*2 gene. The *OCA*2 − *HERC*2 region is thought to play an important role in skin and eye pigmentation in world-wide populations [36].

Next, we analyzed *A. thaliana* data from the 1001 Genomes Project with LODESTAR [20]. We associated each sample with the Cartesian centroid of the sample’s population (Figure 3c) and performed our LODESTAR analysis between non-overlapping windows consisting of 500 loci (Figure 3A). The window with the highest *t*-statistic, *t* = 0.466, is defined by the region *chr*2 : 16719748 − 16770137 (Figure 3b). This window contains the *ELF* 4 gene, which is known to control circadian rhythms and the flowering times of *A. thaliana* [37]. The peak in the *t*-statistic on Chromosome 3 contains *CY P* 72*A* genes. This gene family has been recently tied to seed dormancy [38].

**Figure 3:**
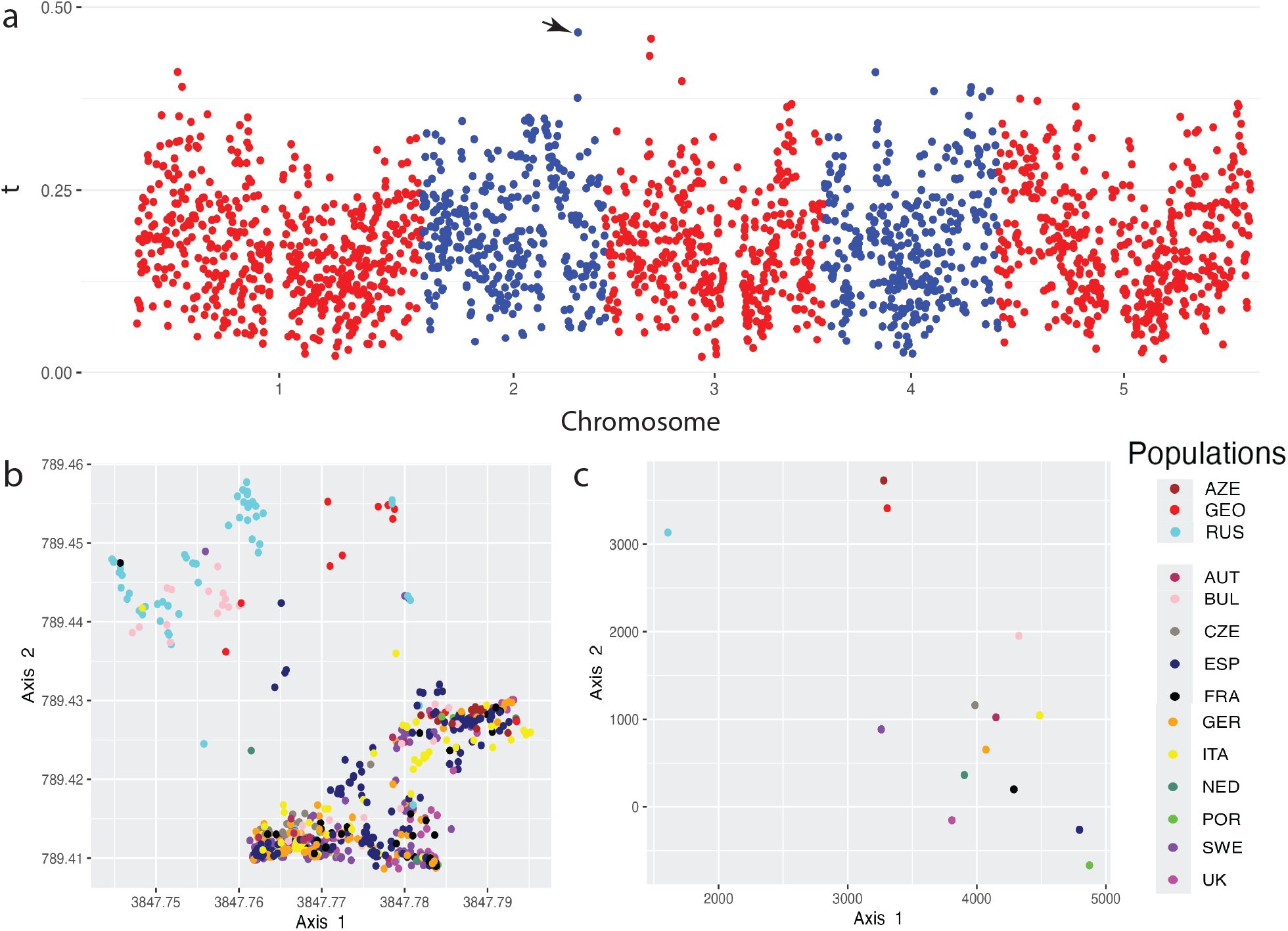
1001 Genomes Project. **Panel a** Procrustes statistic with non-overlapping, 500 loci windows. The window indicated by the arrow is shown in Panel b. **Panel b** The window *chr*2 : 16719748 − 16770137 with the highest Procrustes statistic throughout the genome *t* = 0.466. **Panel c** The centroids of the transformed geographic coordinates in the Cartesian plane.

### 3.2 Genes that Preserve Population Stratification

Often, the geographic locations of samples do not accurately reflect the genetic dissimilarity between samples. Complex demographic histories obscure local patterns of relatedness from the genome-wide realized pattern of relatedness. Locating such regions is informative to explaining the maintained genetic differences between populations when the variation explained by geographic dispersal is limited. We can locate regions that preserve population stratification with LODESTAR by comparing local relatedness plots and the genome-wide relatedness plot during Procrustes analysis. We explored regions of maintained population stratification with all samples from the 1000 Genomes Project, including the trios added at a later date [39]. The total set comprises 3203 samples (Table S1). We ran LODESTAR with non-overlapping, 1000 loci windows (Figure S1). The three highest Procrustes statistics were 0.851136, 0.851290, and 0.853532. The window with *t* = 0.851136 was within *chr*19 : 51036349 − 51291857. To increase our resolution to observe relatedness patterns on Chromosome 19, we performed a LODESTAR analysis using non-overlapping, 500 loci windows (Figure 4a).

**Figure 4:**
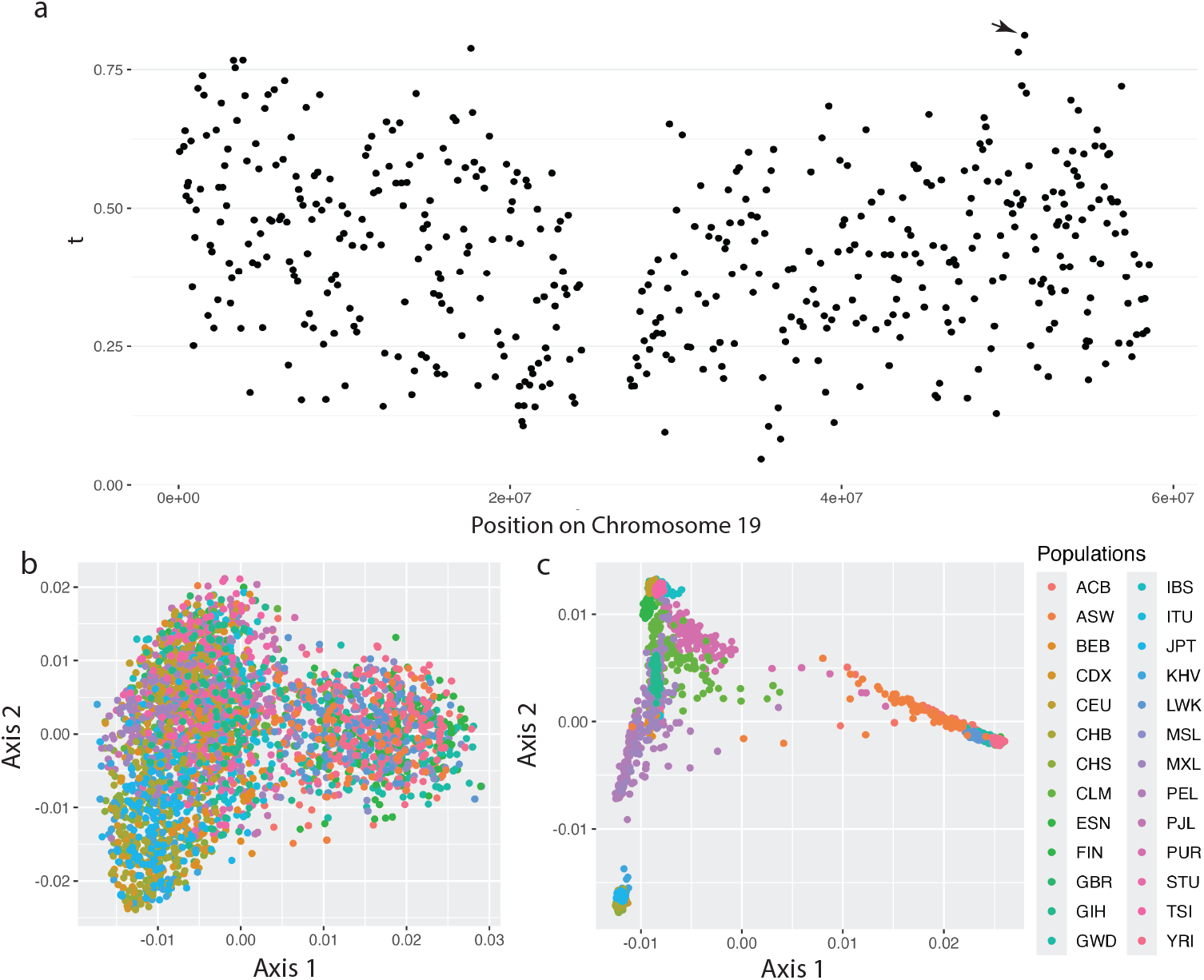
1000 Genomes Chromosome 19. **Panel a** The Procrustes statistic between local, non-overlapping 500 loci windows on Chromosome 19 and the genome-wide relatedness plot for all samples from the 1000 Genomes Project shown in Panel c. The arrow indicated the window shown in Panel b. **Panel b** The window *chr*19 : 51036349 − 51131617 with the highest Procrustes statistic on Chromosome 19 with *t* = 0.811. **Panel c** The genome-wide relatedness plot.

The window (Figure 4b) covers the region *chr*19 : 51036349 − 51131617 and has a Procrustes statistic of *t* = 0.811. The high similarity between this window is seen when compared to the genome-wide plot of relatedness (Figure 4c). Chromosome 19 containing the highest similarity signals to the genome-wide plot of relatedness may not be surprising since it is the most gene-rich chromosome with many duplicated gene families [40]. However, the region (Figure 4b) contains several members of the *KLK* gene family and is directly upstream of the *SIGLEC* genes. Both gene families are related to innate immune function [41, 42]. Immunological diversity is a defining characteristic of differences between populations [43].

### 3.3 Regions Lacking Structure

The change in the Procrustes statistic along the genome can be used as a proxy to observe the change in overall relatedness along the genome. When sampling locations or clear genome-wide population stratification is absent, this change in relatedness is difficult to interpret in an evolutionary context. A more straightforward analysis is to summarize the overall relatedness between samples within the lower-dimensional space. We proposed a measure of this, *σ*, which captures the spread of the samples along the major axes of variation. Therefore, regions of the genome with a high *σ* value are expected to lack population structure, and conversely, regions with a low *σ* maintain population stratification. The *σ* value gives us a measure of distortion in relatedness along the genome. We calculated *σ* in non-overlapping, 1000 loci windows on Chromosome 6 of 99 CEU, 108 YRI, and 103 CHB samples from the 1000 Genomes Project (Figure 5a) [19]. We see a peak in *σ* around the MHC-complex, a known region of immunological genes with diverse alleles in and between populations [44]. The peak next to the MHC-complex in Figure 5a belongs to the centromeres. The window with the highest *σ* defines the region *chr*6 : 33046526 − 33087381 (Figure 5b) and is within the MHC-complex. We juxtapose this region to the window with the lowest *σ* value on Chromosome 6 (Figure 5c). There is a clear difference in the dispersal of samples between the two windows. The window with the highest *σ* lacks structure (Figure 5b) while the window with the lowest *σ* shows separation between the three populations (Figure 5c). Lastly, we plotted the first component, prior to Procrustes analysis, for all samples from the three populations along Chromosome 6 (Figure S2). There is no clear signal indicating high variation between samples within the MHC-complex. This provides evidence that *σ* contains more information compared to the first axis of variation.

**Figure 5:**
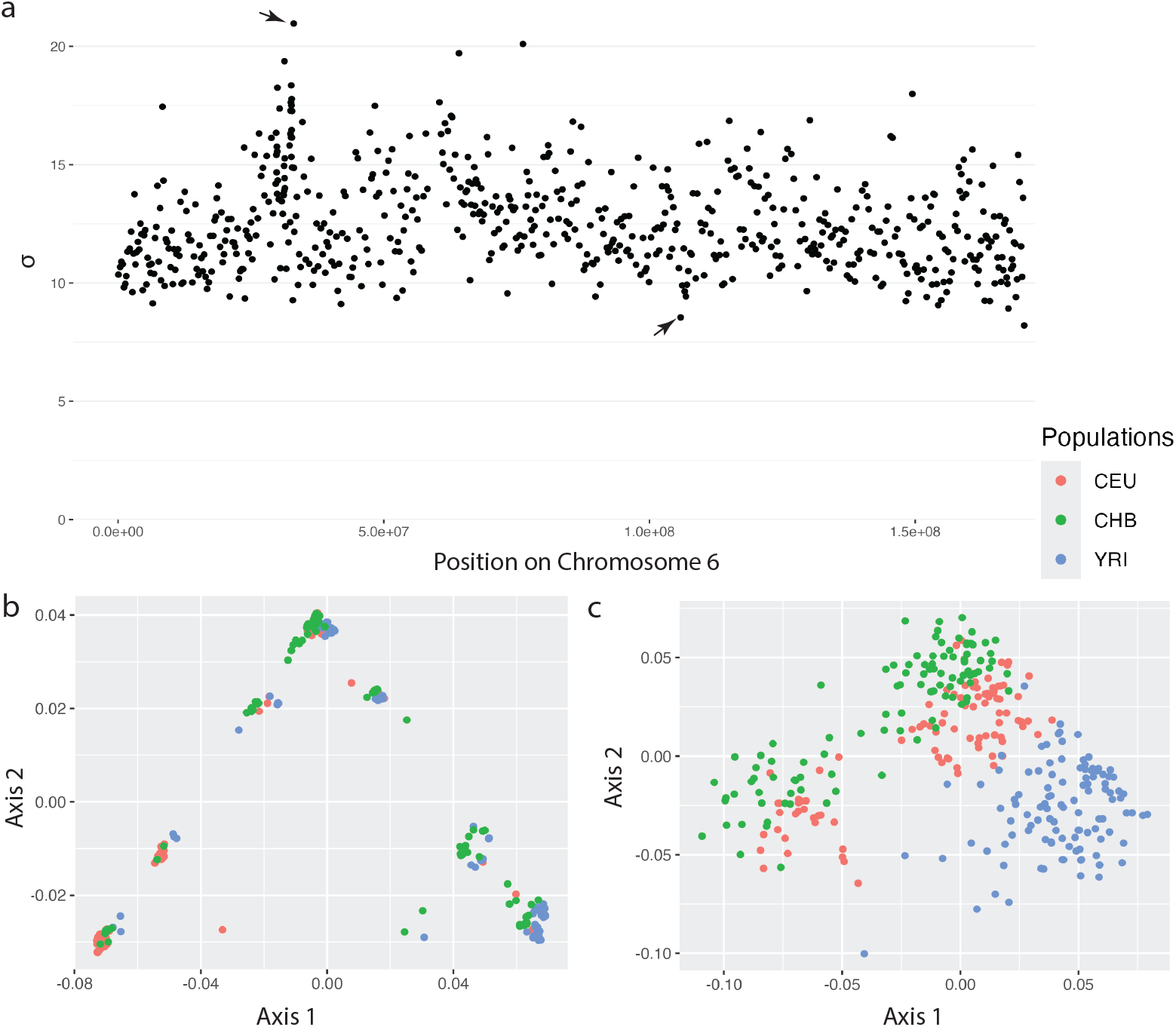
CEU, YRI, CHB Chromosome 6. **Panel a** *σ* in non-overlapping, 1000 loci windows on Chromosome 6 of 99 CEU, 108 YRI, and 103 CHB samples from the 1000 Genomes Project. The left arrow points to the window shown in Panel b. The right arrow points to the window shown in Panel c. **Panel b** The window with the highest value of *σ* = 20.966 on Chromosome 6. **Panel c** The window with the lowest value of *σ* = 8.5466 on Chromosome 6.

### 3.4 Inverted Segments

Recently, PCA has been used to visualize inverted segments. The first principal component for every sample within each window plotted along a genomic region depicts a clear signal associated with an inversion [15, 16]. We illustrate that ASD with cMDS sufficiently captures the inverted segment within a chromosome. Using LODESTAR, we computed the first component of each ST and AR sample within non-overlapping, 100 loci windows along Chromosome 3. We see separation between the ST and AR samples spanning the length of the inverted segment in Figure 6. The inferred differentiation extends beyond the boundaries of the inversion breakpoints into the proximal and distal regions of the inversion.

**Figure 6:**
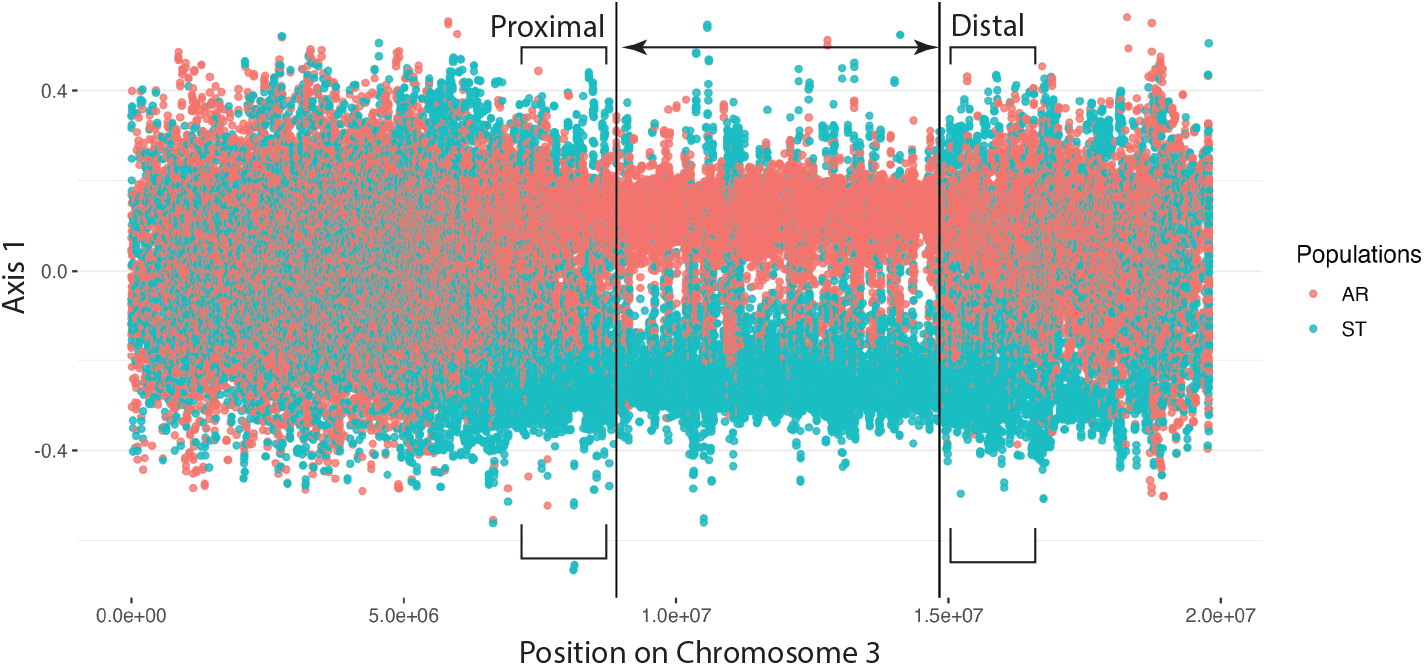
ST and AR Chromosome 3. The first component of each ST and AR sample within non-overlapping, 100 loci windows along Chromosome 3. The Procrustes transformation was not performed. Vertical black bars indicate position of the inversion’s breakpoints.

## 4 Discussion

We presented LODESTAR, a general method to inspect local patterns of genetic relatedness. LODESTAR computes the genetic similarity between samples using pairwise allele-sharing distance. cMDS is used to represent the distances between samples in lower-dimensional space. Then, Procrustes analysis is used to find the optimal rotation and scaling of the local relatedness plot onto a target set of points. Procrustes analysis results in the Procrustes statistic, *t*, which measures the structural correlation between the two sets of points [18]. The target set of points can be the genome-wide relatedness plot or the Cartesian representation of the geographic map of samples. In the case of the target set of points being the genome-wide plot of relatedness, regions with a high Procrustes statistic are regions that preserve population stratification. In the case of the target set of points being a geographic map of samples, regions with a high Procrustes statistic are regions that reflect the sampling geography of the points.

We explored regions that reflected geography in European individuals from 1000 Genomes Project [19]. The region with the highest similarity to the sampling geography was the *OCA*2 − *HERC*2 region. This region plays an important role in skin and eye pigmentation [36], furthering our evidence that human skin and eye pigmentation is deeply tied to human evolution [45]. Then, we located the most similar genomic region in *A. thaliana* from a subset of the 1001 Genomes Project [20] to the samples’ geographic distribution. The *ELF* 4 was in the region with the highest Procrustes statistic. This gene controls the circadian rhythm and flowering times of *A. thaliana*, which are both regulated by temperature, and therefore, are intimately tied to geographic positioning [46]. We performed a LODESTAR analysis using the genome-wide plot of relatedness as a target, including every sample from the 1000 Genomes Project. The genomic region that was most structurally similar to the genome-wide pattern contained *KLK* genes. This gene cluster is responsible for immune related processes and diseases [47]. The genetic mechanisms of the immune system are thought to be key differences between populations and ancestry [48].

Preforming Procrustes analysis along the genome, captures the changes in relatedness between samples. The change in Procrustes statistic could be used as a relative measure for change in relatedness, but we found that a measure of the standard deviation of the samples in lower-dimensional space, *σ*, is more easy to interpret. Regions with high *σ* between samples indicate a high degree of variability. We demonstrated that a region of known diversity, the MHC-complex, shows high variability and lacks population structure [44]. In addition, we showed that ASD in combination with cMDS is sufficient at identifying the position of inversion breakpoints. Similar to the use of PCA to identify inversions, the first axis of variation plotted along Chromosome 3 of *D. pseudoobscura* captured the inverted segment differing between two populations, ST and AR. In addition, differentiation extends beyond the breakpoint boundaries. Recombination suppression within the inverted regions of inversion heterokaryotypes is responsible for differentiation between ST and AR. Differentiation extending into the proximal and distal regions near inversion breakpoints supports the idea that recombination suppression effect in inversion heterozygotes extends beyond the boundaries of the inversion.

We introduced LODESTAR as a qualitative tool to identifying genomic regions of potential evolutionary influence. LODESTAR lacks a rigorous statistical test to evaluate the significance of such identified regions, leading to possible spurious results. Therefore, we recommend further scrutiny of candidate regions identified by LODESTAR within the biological context of the data being studied. We explored the use of a permutation test to evaluate the significance of the Procrustes statistic for each window [25]. The permutation test consisted of shuffling the sample labels in the local window and recalculating the similarity measure to generate a null distribution for the Procrustes statistic. We ultimately found the permutation test to be uninformative; the combined use of ASD with cMDS resulted in structurally dissimilar graphs as the number of samples increased, causing the permutation test to always result in significant *p*-values. This is not unusual since genetic similarity measures struggle with statistical inference [22, 49]. Another approach to measuring statistical significance would be to implement a weighted-jackknife or bootstrap to generate evaluate significance [50, 51].

Another challenge, not unique to LODESTAR but intrinsic to most methods employing windows, is the choice of window size [12]. The optimal window size is dependent on many factors such as density of markers, sequence coverage, and recombination rate [52]. However, we argue that precise window sizes are not necessary since LODESTAR is an exploratory method and individually inspects windows as opposed to viewing the relatedness between windows in aggregate [12]. We recommend starting off with manageable window sizes and incrementally refining the window and step sizes as the user desires. When both parameters are sufficiently tuned, the user can manually inspect the results and continue analyses accordingly.

Several possible extensions to LODESTAR exist. The first extension would be to explore more complex evolutionary scenarios, such as admixture and adaptive introgression. Either scenario may show distinct patterns in *t* or *σ*. Another extension would be to explore the association between local patterns of relatedness and other non-genetic factors. This would shine light on the interaction between genetics and other environmental factors, adding to the conversation on the impacts of climate change [53]. Lastly, LODESTAR can be used to verify recapitulated demographic histories through simulations. Machine learning algorithms are trained on simulated data based on hypothesized demographic histories of real populations [54, 55]. High similarity between the lower-dimensional representation of the simulated samples compared to the realized population structure provides evidence that the simulated demographic history is accurate.

We have proposed a flexible method to inspect the relatedness between samples in local genomic regions and the similarities of this structure to genome-wide relatedness and geographic relatedness. In addition, we described how local genomic relatedness patterns offers a direct interpretation of regions with high diversity and inversions. We demonstrated applications of LODESTAR in each of the situations. This framework could potentially be useful in disentangling evolutionary processes and inform efforts in conversation and evolutionary biology.

## 5 Acknowledgments

Computations for this research were performed using the Pennsylvania State University’s Institute for Computational Data Sciences’ Roar supercomputer. This work was supported by the National Institute of General Medical Sciences of the National Institutes of Health award number R35GM146926 (ZAS), and start-up funds from the Pennsylvania State University’s Department of Biology (ZAS).

## 6 Supplemental

**Table S1:** Populations used from 1000 Genomes Project.

| Number of Samples | Population | Population Code |
| --- | --- | --- |
| 116 | African-Caribbean | ACB |
| 74 | African-American SW | ASW |
| 131 | Bengali | BEB |
| 93 | Dai Chinese | CDX |
| 179 | CEPH | CEU |
| 103 | Han Chinese | CHB |
| 163 | Southern Han Chinese | CHS |
| 132 | Colombian | CLM |
| 149 | Esan | ESN |
| 99 | Finnish | FIN |
| 91 | British | GBR |
| 103 | Gujarati | GIH |
| 178 | Gambian | GWD |
| 157 | Spanish | IBS |
| 107 | Indian | ITU |
| 104 | Japanese | JPT |
| 122 | Kinh Vietnamese | KHV |
| 99 | Luhya | LWK |
| 99 | Mende | MSL |
| 97 | Mexican-American | MXL |
| 122 | Peruvian | PEL |
| 146 | Punjabi | PJL |
| 139 | Puerto Rican | PUR |
| 114 | Sri Lankan | STU |
| 107 | Tuscan | TSI |
| 178 | Yoruba | YRI |

**Figure S1:**
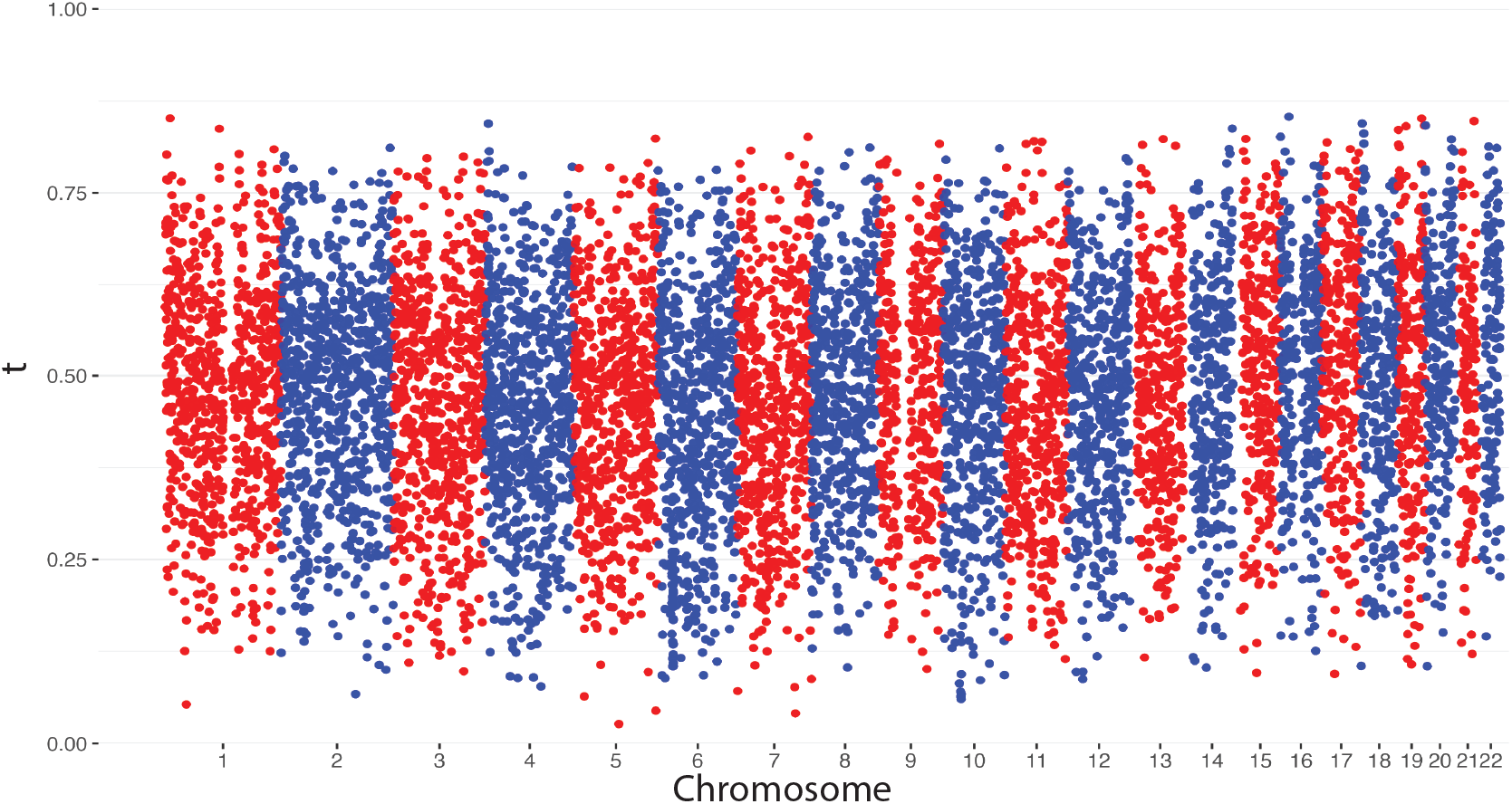
1000 Genomes LODESTAR Analysis. The Procrustes statistic between local, non-overlapping 1000 loci windows and the genome-wide relatedness plot for all samples from the 1000 Genomes Project.

**Figure S2:**
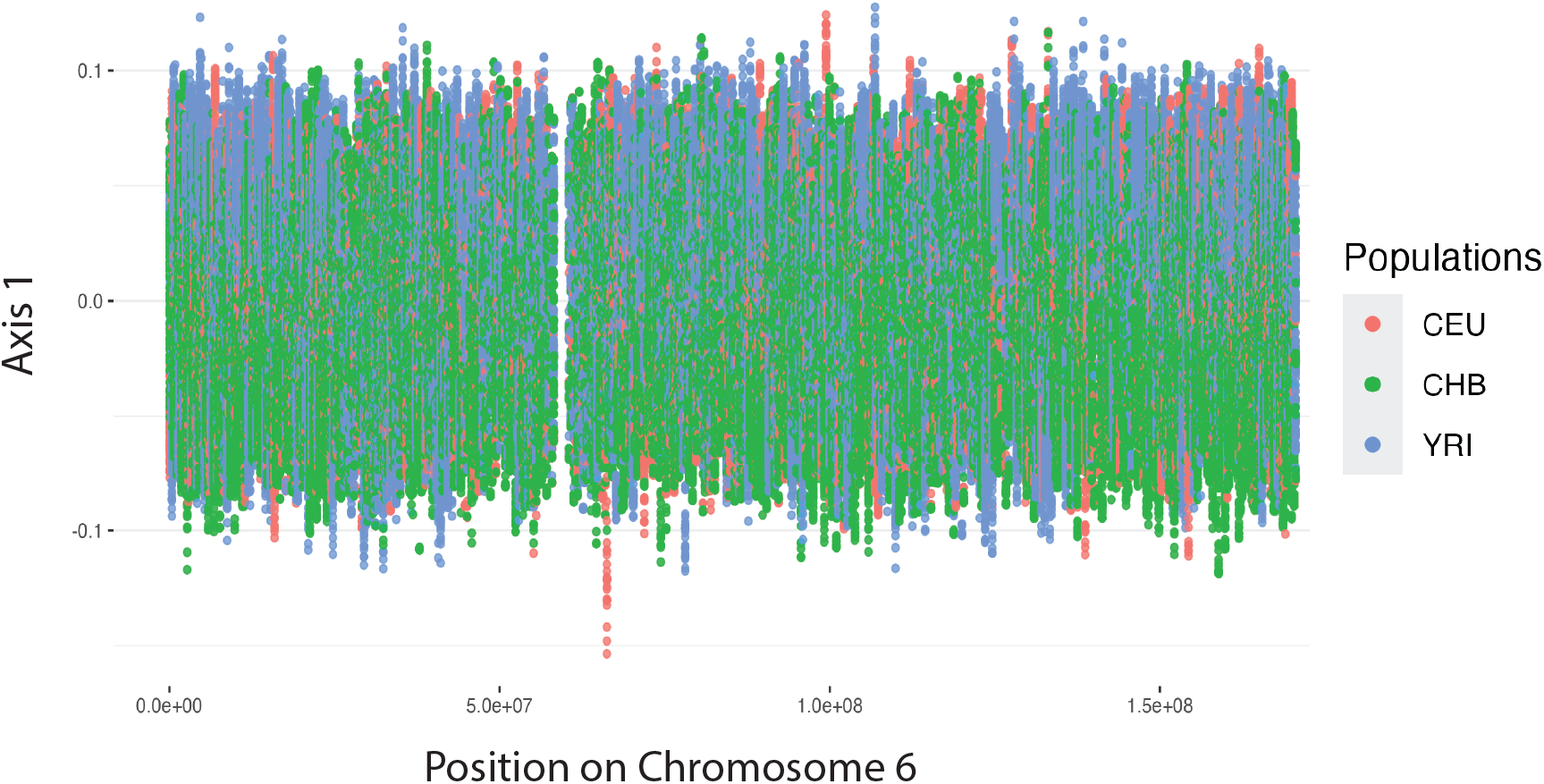
First Component of CEU, YRI, CHB Chromosome 6. The first component, prior to Procrustes analysis, in non-overlapping, 1000 loci windows on Chromosome 6 of 99 CEU, 108 YRI, and 103 CHB samples from the 1000 Genomes Project.

**Table S2:** Populations used from 1001 Genomes Project.

| Number of Samples | Population | Population Code |
| --- | --- | --- |
| 13 | Austria | AUT |
| 16 | Azerbaijan | AZE |
| 28 | Bulgaria | BUL |
| 39 | Czechia | CZE |
| 180 | Spain | ESP |
| 45 | France | FRA |
| 14 | Georgia | GEO |
| 118 | Germany | GER |
| 73 | Italy | ITA |
| 11 | Netherlands | NED |
| 10 | Portugal | POR |
| 60 | Russia | RUS |
| 243 | Sweden | SWE |
| 69 | United Kingdom | UK |

